# Multi-resolution characterization of molecular taxonomies in bulk and single-cell transcriptomics data

**DOI:** 10.1101/2020.11.05.370197

**Authors:** Eric R. Reed, Stefano Monti

## Abstract

As high-throughput genomics assays become more efficient and cost effective, their utilization has become standard in large-scale biomedical projects. These studies are often explorative, in that relationships between samples are not explicitly defined *a priori*, but rather emerge from data-driven discovery and annotation of molecular subtypes, thereby informing hypotheses and independent evaluation. Here, we present *K2Taxonomer*, a novel unsupervised recursive partitioning algorithm and associated R package that utilize ensemble learning to identify robust subgroups in a “taxonomy-like” structure (https://github.com/montilab/K2Taxonomer). *K2Taxonomer* was devised to accommodate different data paradigms, and is suitable for the analysis of both bulk and single-cell transcriptomics data. For each of these data types, we demonstrate the power of *K2Taxonomer* to discover known relationships in both simulated and human tissue data. We conclude with a practical application on breast cancer tumor infiltrating lymphocyte (TIL) single-cell profiles, in which we identified co-expression of translational machinery genes as a dominant transcriptional program shared by T cells subtypes, associated with better prognosis in breast cancer tissue bulk expression data.

## Introduction

As high-throughput transcriptomic assays become more efficient and cost-effective, they are being routinely integrated into large-scale biomedical projects^1–4^. Bulk gene expression profiling by RNA sequencing (RNAseq) has been widely adopted in multiple high-throughput genomics studies, the paramount example being The Cancer Genome Atlas (TCGA) data commons, which currently include 10,558 bulk RNA sequencing (RNAseq) profiles across 33 cancer types (https://portal.gdc.cancer.gov/). Furthermore, since its first published application in 2009^5^, the size of single-cell RNA sequencing (scRNAseq) studies has exploded, such that is now commonplace for studies to generate tens of thousands of profiles^6^. As the scale of these studies and the associated datasets increases, so does their utility as a resource from which biological information can be extracted through the application of machine learning approaches. Common deliverables of these types of analysis include the discovery and characterization of molecular subtypes, which are prevalent in both bulk and single-cell gene expression studies. For example, TCGA bulk expression data have been utilized to characterize subtypes of numerous cancers^7^, including but not limited to: breast^8^, colorectal^9^, liver^10^, and bladder cancer^11,12^. Similarly, the characterization of molecular subtypes is a standard component of the scRNAseq data analysis workflow, insofar as estimation and annotation of subpopulations of cells is one of the primary goals of the assay^13^.

The general framework for subtype characterization can be summarized in two steps: 1) estimation of data-driven groups of observations via application of an unsupervised learning procedure, followed by 2) annotation of each group based on the identification of distinct patterns of gene expression relative to other groups. While most approaches focus on discovering a “flat” set of non-overlapping groups or subtypes, in this manuscript we present an alternative approach, devised to emphasize “taxonomy-like” hierarchical relationships between observations to discover nested subgroups.

Whereas a wide range of unsupervised learning algorithms is available for the analysis of bulk gene expression data, the considerable sparsity of scRNAseq data has motivated the development of novel methods specifically tailored to the analysis of this type of sparse, high-dimensional data. Popular software packages, such as Seurat^14^ and Scran^15^, generate “flat” clusters, in which a finite set of mutually exclusive cell types is estimated. In so doing, they fail to capture the “taxonomy-like”, hierarchical structure that may exist among subgroups of observations at multiple levels of resolution, driven by transcriptional signatures based on different factors, including but not limited to: shared lineage, cell state, pathway activity, or morphological origin. Complementary methods exist to model such relationships, such as *Neighborhood Joining*^16,17^ and more recent single-cell *trajectory inference* approaches^18^, which estimate “pseudo-temporal” states of individual cells indicative of developmental progression. Given the stringent interpretation of such models, their suitability depends on the assumption that the measured similarity between neighbors of cell profiles arises from a distinct continuous progression of molecular activity. However, the relative similarity between cell profiles may be confounded by numerous factors, including: cell cycle, spatial patterning, and cell stress, and batch effects^19^. To overcome these shortcomings, one recent method, partition-based graph abstraction (PAGA)^20^, was devised to model complex topologies by estimating a graph of “high-confidence” connections between labeled cell types based on their shared nearest neighbors. This method has the advantage of being able to first identify disconnected subgraphs from which to model separate trajectories. Even so, a ubiquitous step of *trajectory inference* approaches, including PAGA, is that all distances between cell profiles are computed based on a single set of features, generated by selection filtering and/or dimensionality reduction^21^, thus precluding the discovery of nested structures defined by distinct transcriptional programs shared by relatively few cells.

Hierarchical clustering (HC) algorithms at face value address the need for a multi-resolution representation of the relationship among observations, and while originally adopted for the analysis of bulk gene expression data^22^, numerous packages have also been developed for scRNAseq analysis, such as *pcaReduce*^23^, *ascend*^24^, and *BackSPIN*^25^. However, since the number of possible subgroupings increases with the number of observations, robustly identifying such relationships can be challenging. As a result, tree-cutting methods are often applied, ultimately yielding a flat set of non-overlapping clusters. Furthermore, as with *trajectory inference*, the bottom-up nature of HC’s sample aggregation procedure forces the use of the same set of genes/features to drive the agglomeration at all levels of the hierarchy.

Here we introduce *K2Taxonomer*, a novel taxonomy discovery approach and associated R package for the estimation and in-silico characterization of hierarchical subgroup structures in both bulk and single-cell data. An important feature of the approach is that it can analyze both individual samples as well as sample groups such as, but not limited to, those corresponding to scRNAseq cell types. The package employs a recursive partitioning algorithm, which utilizes repeated perturbations of the data at each partition to estimate ensemble-based K=2 subgroups. For scRNAseq analysis, *K2Taxonomer* utilizes the constrained k-means algorithm^26^, to estimate partitions of the data at the cell type level, while preserving the influence of each individual cell profile. A defining feature of the method is that <u>each recursive split of the input data is based on a distinct set of features</u> selected to be most discriminatory within the subset of samples member of the current hierarchy branch. This makes the approach quite distinct from a standard clustering algorithm, and particularly apt to discover nested taxonomies. In addition, the package includes functionalities to comprehensively characterize and statistically test each subgroup based on their estimated stability, gene expression profiles, and a priori phenotypic annotation of individual profiles. Importantly, all results are aggregated into an automatically generated interactive portal to assist in parsing the results.

In this manuscript, we assess the performance of *K2Taxonomer* for partitioning both bulk gene expression and scRNAseq data, using both simulated and publicly available data sets, and we compare it to agglomerative clustering procedures. For bulk gene expression data, performance is assessed in terms of unsupervised sorting of breast cancer subtypes and established genotypic markers, using breast cancer patient tumor tissue data from the Molecular Taxonomy of Breast Cancer International Consortium (METABRIC)^27^ and the TCGA compendia.

For scRNAseq data, performance is assessed in terms of recapitulation of established relationships between 28 annotated cell types of the airway of healthy subjects^28^. We conclude with a case study where we perform a *K2Taxonomer*-based analysis of breast cancer tumor infiltrating lymphocytes (TILs) profiled by scRNAseq^29^. Our analysis significantly expands upon previously published results and identifies a phenotypically diverse subgroup of CD4 and CD8 cells, characterized by constitutive up-regulation of a subset of translation machinery genes. We further show that high expression of these genes in breast cancer tissue bulk expression is associated with better survival, supporting recent findings on the role of the translation machinery assembly in T cell activation^30,31^, and demonstrate that this coordinated expression of the translation machinery is pervasive among T cell subpopulations to such an extent that the expression levels of these genes in bulk measurements of tumor tissue is predictive of the degree of immune infiltration. The complete suite of analysis results is accessible through an automatically generated and publicly accessible portal (https://montilab.bu.edu/k2BRCAtcell/).

While we focus on the analysis of transcriptomics data, we emphasize that our approach is applicable to other ‘omics’ data, such as those generated by high-throughput proteomics and metabolomics assays. Moreover, applications of *K2Taxonomer* are not limited to cancer-centric studies. For example, we applied an earlier prototype of *K2Taxonomer* to the analysis of a toxicogenomic study on the effects of environmental exposures on adipocyte activity, and the tool proved to be instrumental to the identification of chemical subgroups and the pathways contributing to either their deleterious or beneficial effects on energy homeostasis^32^.

## Results

### K2Taxonomer discovers hierarchical taxonomies on simulated data

We first evaluated *K2Taxonomer*’s capability to recapitulate hierarchical relationships induced in simulated data, as measured by the Baker’s gamma coefficient estimate of similarity between the structure of two dendrograms, where the structure of each dendrogram is quantified by a matrix based on the number of partitions separating each pair of leaves^33^. We evaluated the method’s performance both on data where the analysis end-points were single samples (*observation-level*), and groups of samples (*group-level*). The latter corresponds to scenarios where the goal is to define a taxonomy over sample groups, such as cell types in single-cell experiments, or chemical perturbations profiled in multiple replicates^34^. As a term of reference, we compared *K2Taxonomer*’s performance to Ward’s agglomerative method.

For observation-level analysis, *K2Taxonomer* demonstrated robust performance for moderate levels of background noise, e.g. standard deviations equal to 0.5 and 1.0, regardless of the proportion of features with signal or the number of terminal clusters (Figure 2a, Table S1). For higher levels of background noise, e.g., standard deviations equal to 2.0 and 3.0, the performance of *K2Taxonomer* was more dependent on either parameter, performing better with more features with signal and fewer terminal clusters. *K2Taxonomer* significantly outperformed Ward’s method in 221 out of the 400 combinations of parameters tested (FDR < 0.05), while Ward’s performed better for 17 combinations (Figure 2a, Table S1). Furthermore, the differences between Baker’s gamma coefficients for the 221 results significantly in favor of *K2Taxonomer* were generally larger than the 17 results significantly in favor of Ward’s method, with median differences of 0.14 and 0.03, respectively. In general, *K2Taxonomer* significantly outperformed Ward’s method when the background noise, the number of terminal groups, and percent features with signal increased. Remarkably, for group-level analysis *K2Taxonomer* outperformed Ward’s method for all 400 combinations of variables tested (Figure 2b, Table S3).

**Figure 1:**
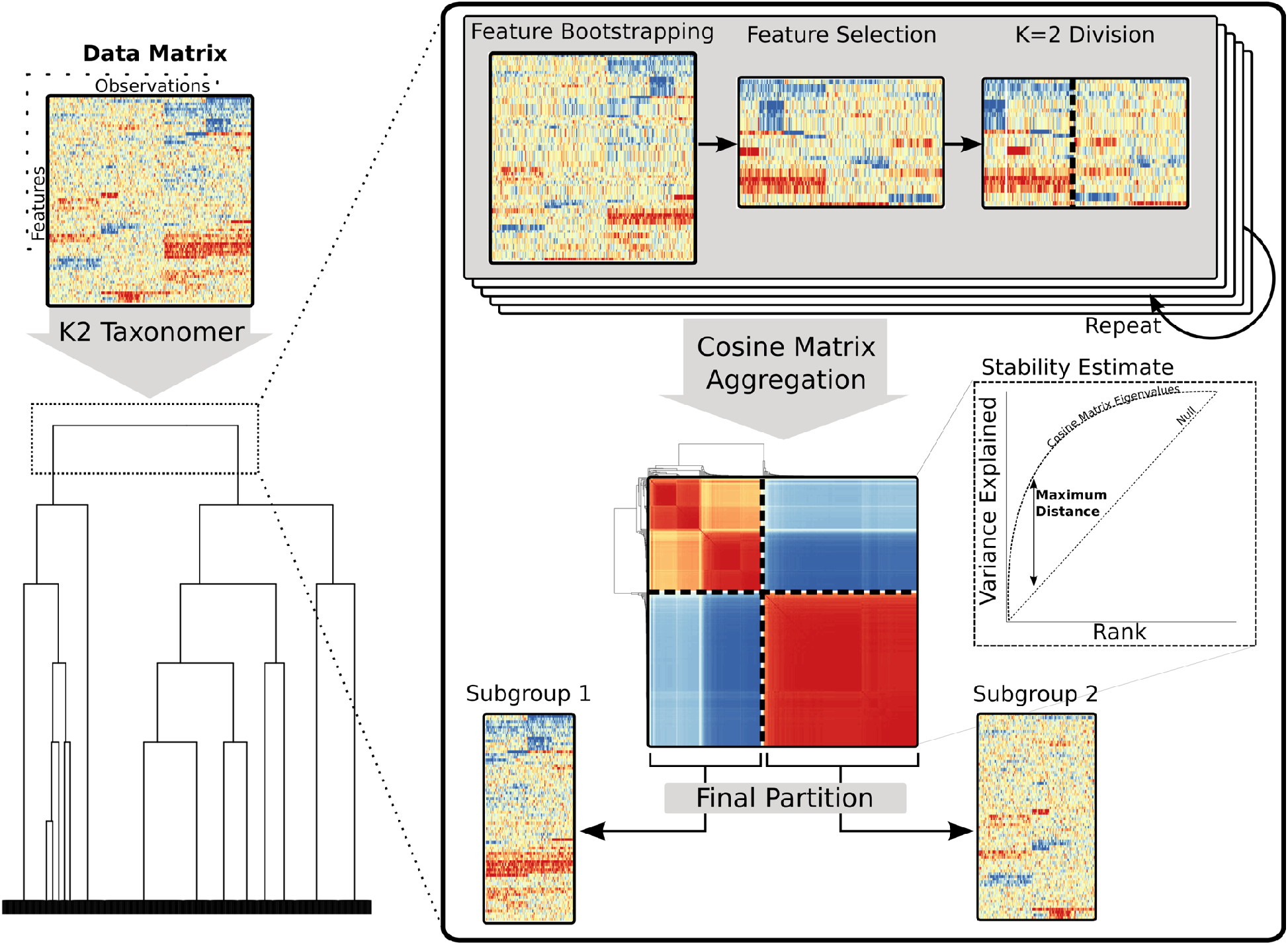
Schematic of the *K2Taxonomer* recursive partitioning algorithm. For each partition, *K2Taxonomer* generates an ensemble of K=2 estimates from the feature bootstrapped data followed by variability-based feature selection. This ensemble is aggregated to a cosine matrix followed by hierarchical clustering and tree cutting. A stability estimate, indicative of the consistency of K=2 estimates, is calculated based on an eigendecomposition of the cosine matrix. *See supplementary methods for a more thorough description of the elements of this procedure.*

**Figure 2:**
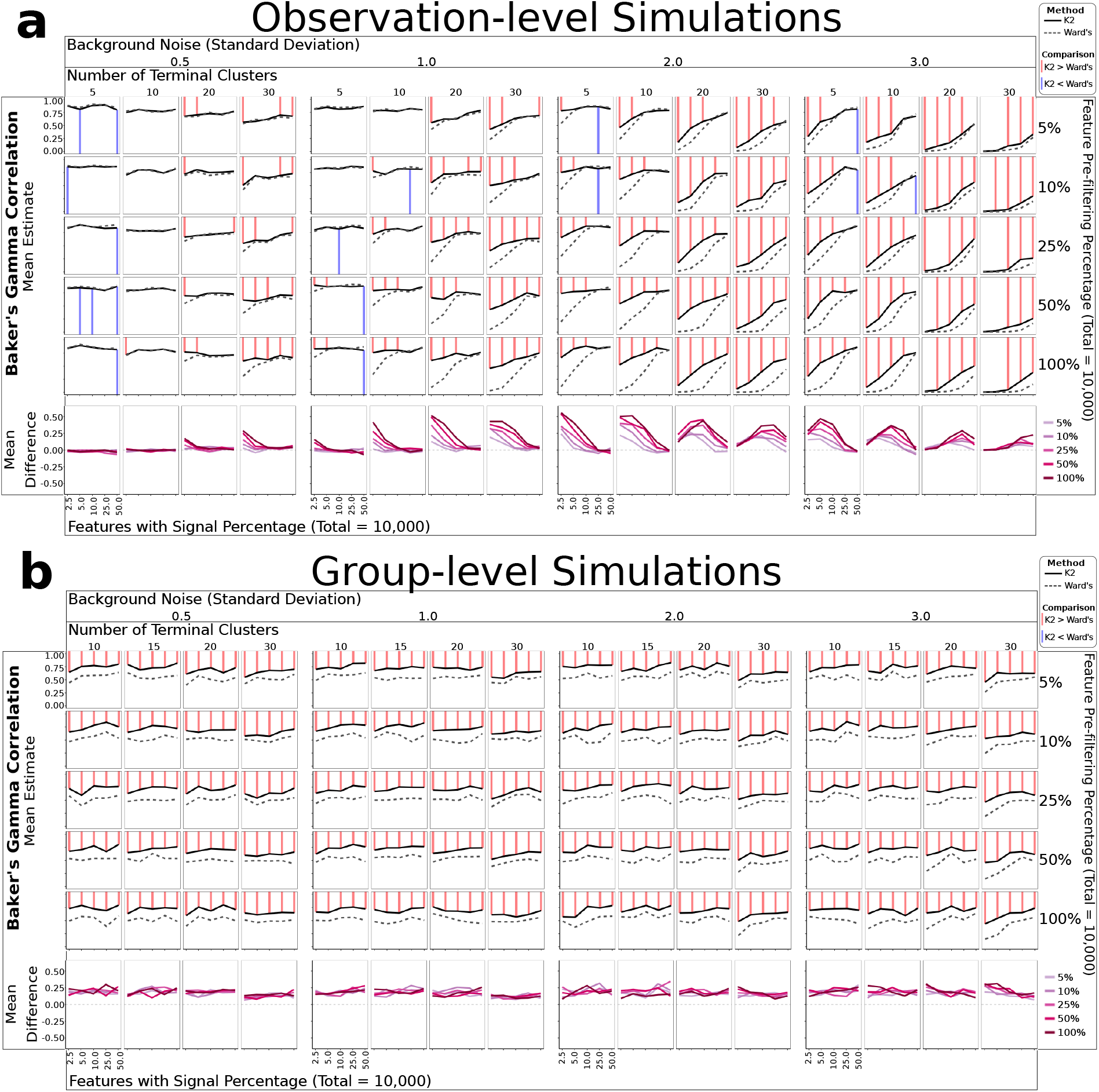
Simulation-based performance assessment of *K2Taxonomer* and Ward’s agglomerative method. Mean Baker’s gamma correlation estimates measuring the similarity of either *K2Taxonomer* and Ward’s agglomerative method estimates to the true hierarchy from which the simulated data was generated. Each combination of parameters was simulated 25 times. The red and blue lines are indicative of statistically significant differences between the correlation estimates (FDR < 0.05) based on a Wilcoxon signed-rank test. A) Observation-level analyses with 300 observations and 10,000 features. B) Cohort-level analyses with 1,000 observations and 10,000 features.

Using the square root of the total number of features as the partition-specific feature filtering parameter for running *K2Taxonomer* demonstrated stable performance. When compared to selecting a fixed percentage of the total number of features (Figure S1, Table S2), the square root outperformed larger percentages when the number of features was large, and outperformed smaller percentages when the number of features was small.

### K2Taxonomer accurately sorts breast cancer subtypes without pre-filtering of features

We evaluated *K2Taxonomer*‘s ability to sort Pam50 subtypes, ER-status, PR-status, and HER2-status from bulk gene expression data from METABRIC and the TCGA BRCA bulk gene expression data, separately. A fourth variable, defined by the Cartesian product of ER-status, PR-status, and HER2-status was also assessed. Performance was assessed in terms of the decrease in entropy as the number of cluster estimates, K, increased from 2 to 8 (Figure 3a-b). We also compared *K2Taxonomer*’s performance to two agglomerative clustering methods, Ward’s and average. Since standard hierarchical clustering is sensitive to the level of feature filtering, the comparison was repeated for multiple pre-filtering levels.

**Figure 3:**
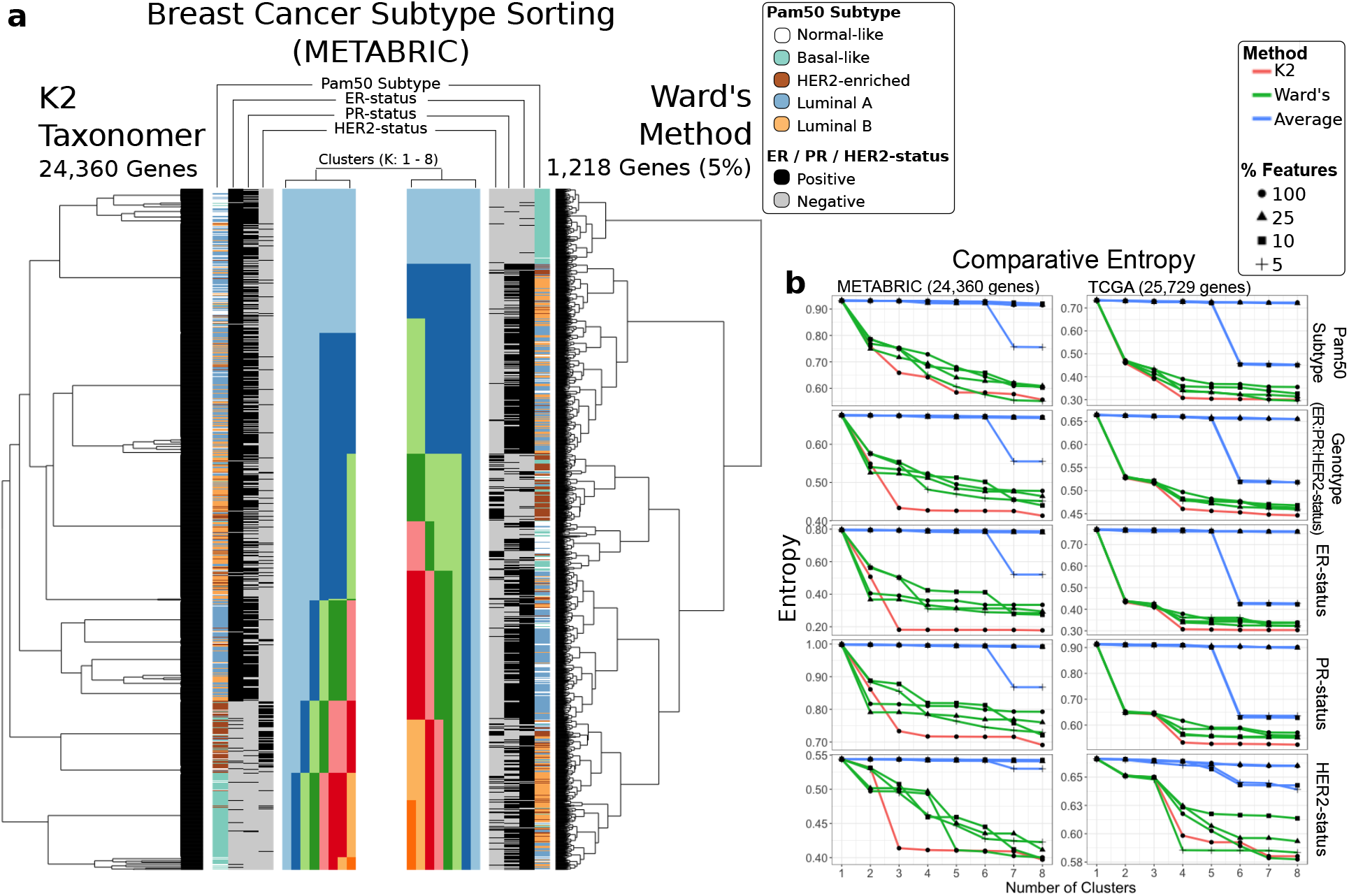
Breast cancer subtyping performance assessment of bulk gene expression data. Comparison of sorting of breast cancer Pam50 subtypes and genotypes (ER-, PR-, and ER-status) for two bulk gene expression data sets, *METABRIC* and *TCGA*.An aggregate, three gene genotype status was also included by combining the individual genotypes. Performance was assessed based on reduction of entropy as the number cluster estimate increased based on tree cutting. *K2Taxonomer* was only run on the full set of features, while either agglomerative method, average and Ward’s, were run on three additional subset of the data. A) Illustration of the results generated by *K2Taxonomer* and Ward’s method for the *METABRIC* dataset. These results reflect Ward’s method run on 5% of the total number of features, which demonstrated the best performance among agglomerative methods. B) Entropy measurements for each method as K increased across the *METABRIC* and *TCGA* data sets.

In general, *K2Taxonomer* accurately segregated the known subtypes and phenotypes, performing as well or better than either method (Figure 3b, Table S6, Table S7). When applied to the METABRIC data, *K2Taxonomer* analysis yielded the lowest entropy score compared to all other methods for K=3 and higher with few exceptions. Other methods produced similar entropy measurements at selected higher levels of K. For example, Ward’s method resulted in similar entropy scores for Pam50 subtypes and HER2-status at K=4 and K=5, respectively, but for different pre-filtering levels, 5% and 100%, respectively. When applied to the TCGA BRCA data, the difference in performance was less pronounced. *K2Taxonomer* resulted in the lowest entropy score for Pam50 scores, genotype, ER-, and PR-status for K=4. Ward’s method at 5% pre-filtering level produced the smallest entropy score for HER2-status for K=4.

It should be emphasized that the pre-filtering level to be used with hierarchical clustering is not known a priori, and it would thus preclude us in practice from selecting the level yielding the best results shown in the above comparison.

### K2Taxonomer accurately identifies and organizes subgroups of shared progenitors and epithelial cells from healthy airway scRNAseq cell clusters

To assess the capability of *K2Taxonomer* to recapitulate biologically relevant subgroupings of cell types estimated from scRNAseq data, we ran group-level analysis using 29 cell types estimations assigned to 77,969 cells of airway tissue from 35 samples across 10 healthy subjects and multiple locations (Figure 4a, b)^24^. In addition to agglomerative methods, we evaluated *K2Taxonomer’s* performance against that of partition*-based graph abstraction (PAGA)*^20^, which has been shown to outperform similar methods, especially for analyzing large-scale scRNAseq data sets^18^.

**Figure 4:**
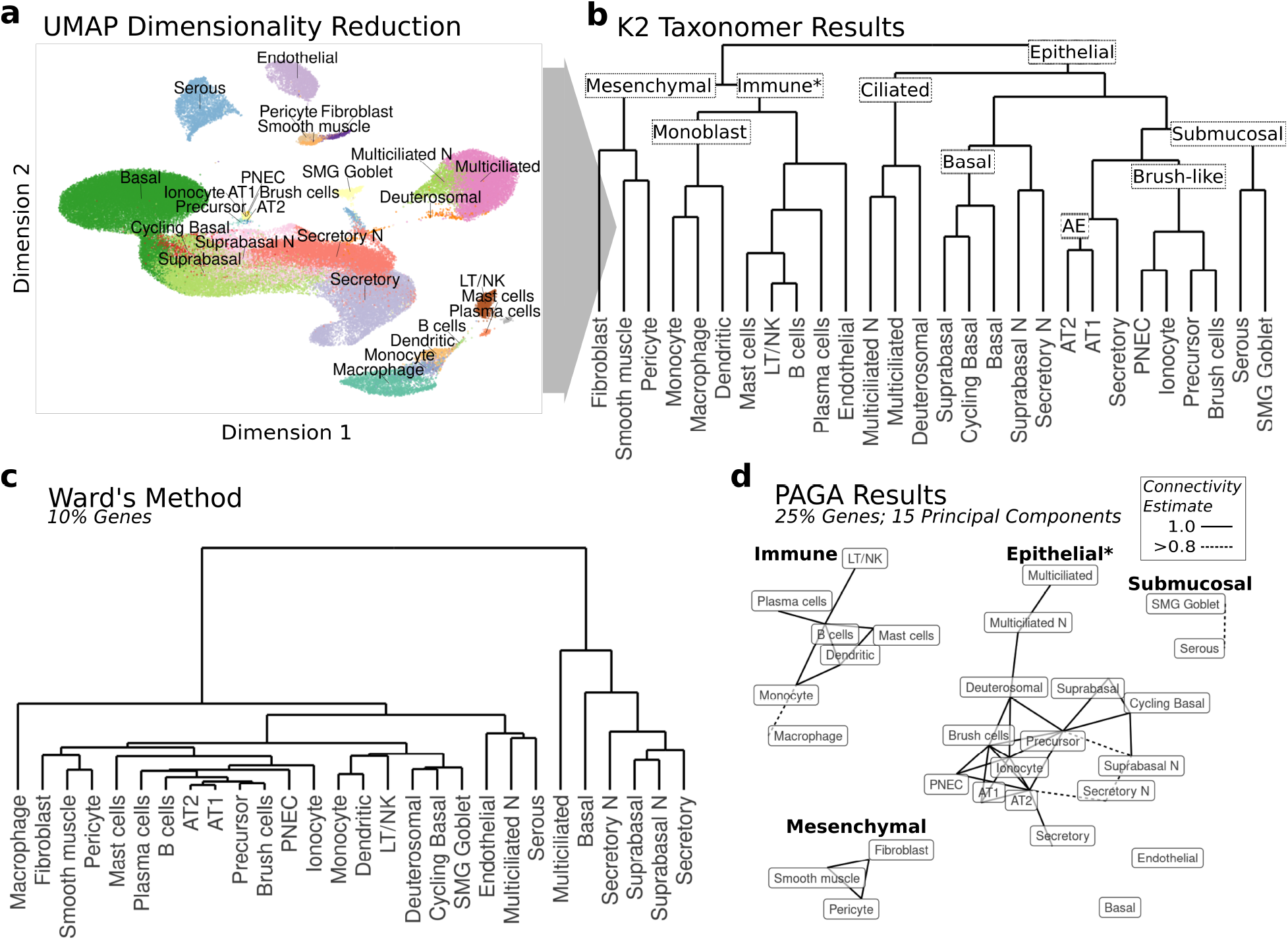
Subgrouping of healthy airway cell types from scRNAseq data. A) tSNE dimensionality reduction of healthy airway scRNAseq data with labels for 28 cell types annotated by ^28^. Note cell types labelled ending in “N” indicate those which only included cells from nasal samples. B) *K2Taxonomer* results with nine annotated subgroups. The “*” in the “Immune” subgroup label indicates the impurity of this subgroup caused by the presence of endothelial cells. C) Ward’s agglomerative clustering results for a selected analysis performed on 10% of the total number of genes. The results for both Ward’s method and Average method run on additional gene subsets: 100%, 25%, 10%, and 5%, are shown in Figure S2. D) PAGA graph-based trajectory results for a selected analysis performed on 15 principal components estimated from 25% of the total number of genes with four annotated disconnected subgraphs. The “*” in the “Epithelial” subgroup label indicates the incompleteness of this subgraph caused by the absence of basal cells and submucosal cells. Edges indicate PAGA connectivity estimates > 0.8. The results for analyses run on additional numbers of principal components: 10, 15, and 20, as well as additional gene subsets: 100%, 25%, 10%, and 5%, are shown in Figure S3.

*K2Taxonomer* was remarkably accurate in capturing the higher-order organization of the 28 cell types. The first partition separated all epithelial cell subtypes from non-epithelial cell types (Figure 4b). Further partitioning of the 17 epithelial cell subtypes yielded five subgroups, characterized by shared morphology, labeled as “Ciliated”, “Basal”, “Submucosal”, “Brush-like”, and “AE” (Alveolar Epithelium). The “Ciliated” subgroup was comprised of differentiated multiciliated cells and their precursor, deuterosomal cells^28^. The “Basal” subgroup was comprised of epithelial cell progenitors, basal and cycling basal cells, as well as epithelial cell intermediary, suprabasal cells^35^. The “Brush-like” subgroup was comprised of three rare airway epithelial cell types: brush, ionocyte, and pulmonary neuroendocrine cells (PNECs), as well as their likely shared “brush-like” precursor cells^36^, as suggested in the original publication of these data^28^. Further partitioning of the 11 non-epithelial cell types yielded two main subgroups characterized by shared progenitor cells: immune^37^ and mesenchymal stem cells^38^, with the immune cell subgroup also including endothelial cells. Further partitions of the immune cell subgroup included progeny of monoblasts: monocytes, dendritic cells, and macrophages^39^, followed by endothelial cells separated from all non-monoblast progeny immune cells subtypes in the adjacent subgroup.

In contrast, agglomerative hierarchical clustering of these cell clusters, even if evaluated at multiple F-statistic-based pre-filtering levels, yielded significantly different results poorly reflective of the known taxonomic cell type organization (Figure 4c, Figure S2). While the mesenchymal stem cell subgroup, comprised of fibroblasts, smooth muscle, and pericytes, was identified by Ward’s method, and while there were other instances of concordant subgroups, none of these consisted of more than two cell types.

*PAGA-based* trajectory analysis^20^ of the cell performed better than agglomerative clustering, and demonstrated both improvements and drawbacks compared to *K2Taxonomer*. (Figure 4d, Figure S3). Figure 4d shows what we selected to be the “best” PAGA result for runs on a grid of F-statistic-based pre-filtering levels (25% of genes) and principal component-based dimensionality reduction size (15 principal components). In this case, *PAGA* recapitulated four subgroups identified by *K2Taxonomer* as disconnected subgraphs: “Immune”, “Epithelial”, “Submucosal”, and “Mesenchymal”. Unlike *K2Taxonomer*, this model accurately segregated endothelial cells as its own disconnected vertex. However, all *PAGA* models failed to include basal cells in the epithelial cell subgraph (Figure 4d, Figure S3). Furthermore, PAGA segregated the histologically separated submucosal cells lines: SMG goblet and serous, from the other epithelial cell lines. The most apparent difference between the *K2Taxonomer* and *PAGA* results is the high connectedness of the PAGA subgraphs, especially considering that these are all “near perfect-confidence” connections. As a result, subgroup relationships within these subgraphs are difficult to distinguish, and approaches to estimate tree-like graphs, such as minimum spanning tree algorithms, yield multiple equivalent solutions. Finally, the PAGA results varied based on the feature pre-filtering level and number of principal components, most notably in the estimations of connections between immune cells and epithelial cells (Figure S3).

### K2Taxonomer identifies subgroups of TILs characterized by differential regulation of TNF signaling, translation, and mitotic activity from BRCA tumor scRNAseq cell clusters

We performed *K2Taxonomer* analysis on scRNAseq data of 13 TIL cell clusters reflecting further subdivision of the 10 cell types reported in the original study^29^. The higher resolution was achieved by reproducing the reported methods^29^ with the exception of selecting a higher resolution parameter when performing clustering with Seurat^14^ (Table S8).

The results of *K2Taxonomer* partitioning and annotation of breast cancer TIL cell clusters estimated from scRNAseq data is summarized in Figure 5a. Biologically informative subgroups, characterized by strongly significant differential expression of gene expression and sample-level pathway enrichment are highlighted and labeled within each boxed sub-dendrogram. The full set of differential results for genes and pathways across all partitions are reported in Table S9 and Table S10, respectively. Three distinct multi-cell subgroups emerged, labeled as: “Trm All”, “CD4+ CCL5−“, and “Translation+”, characterized by consistent up-regulation of PD-1 signaling (Reactome PD-1 signaling, FDR = 1.1e-241), translation (Reactome eukaryotic translation initiation, FDR = 5.6e-137), and TNF signaling (Reactome TNFS bind their physiological receptors, FDR ~ 0.00), respectively (Figure 5b). “Trm All” *and* “Treg” subgroups each included a mitotic cell subgroup characterized by high cell cycle activity (Figure 5B). Furthermore, the “CD4+ CCL5−” subgroup, comprised of the “CD4+ CXCL13+” cell cluster and “Treg” subgroup, is characterized by consistent down-regulation of CCL5 (FDR ~ 0.00) and up-regulation of TNFRSF4 (FDR ~ 0.00) (Figure 5c, d). Furthermore, additional up-regulation of TNFRSF4 (FDR = 1.1e-7) and RGS1 (FDR = 4.3e-55) distinguish non-mitotic “Treg” subgroups (Figure 5c). Gene-level markers of the “Translation+” subgroup included numerous ribosomal proteins, epitomized by up-regulation of RPS27 (FDR = 9.6e-246) (Figure 5c, d).

**Figure 5:**
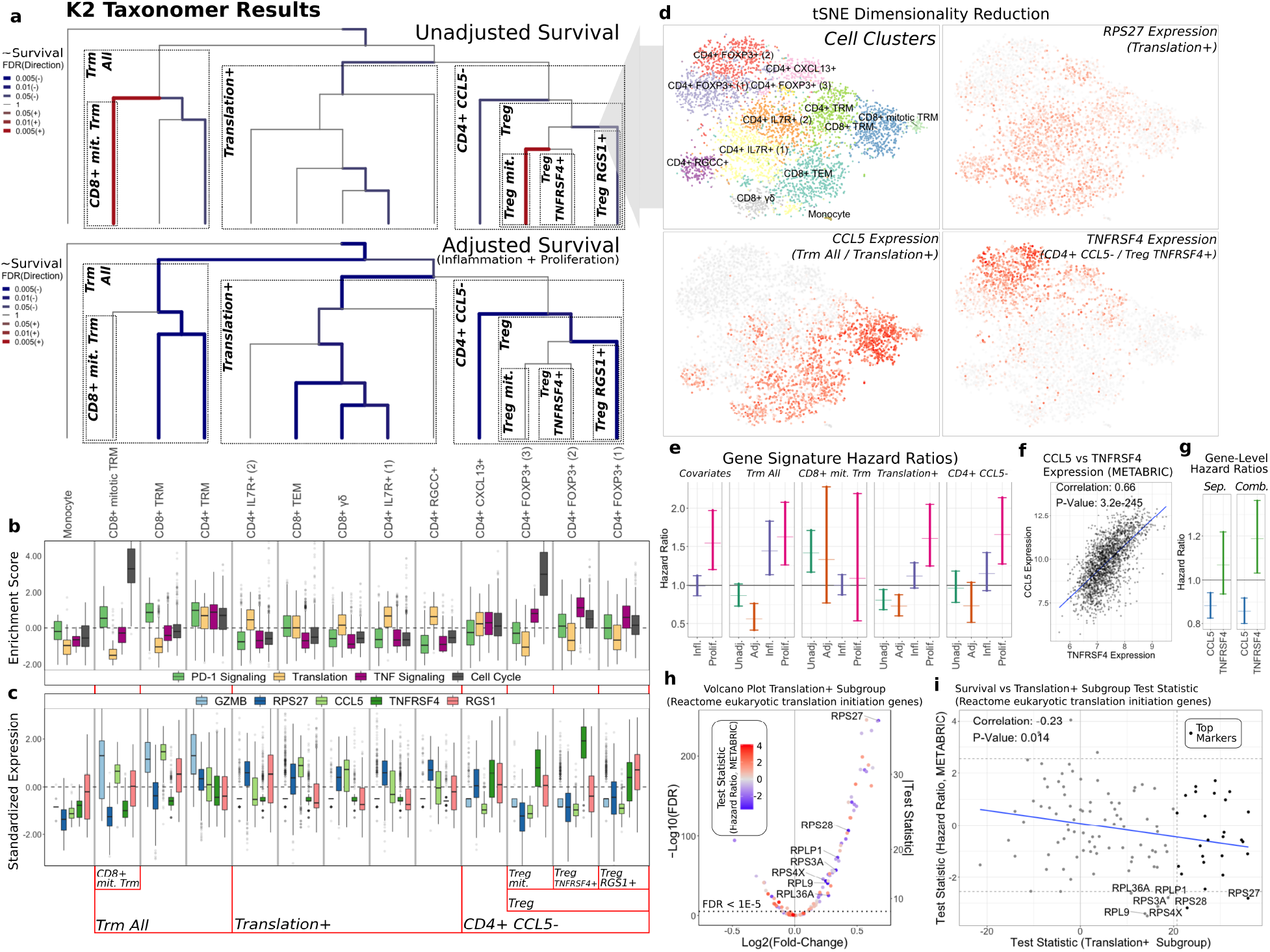
*K2Taxonomer* annotation of scRNA-seq clustering of breast cancer immune cell data and in-silico validation via patient survival on *METABRIC* breast cancer bulk gene expression data set. A) *K2Taxonomer* annotation of 13 cell subtypes of breast cancer immune cell populations. Cell type labels are in accordance to the original publication of these data^29^. Color and thickness of each edge indicates the association between the projected signature of up-regulated genes of each subgroup and patient survival in *METABRIC* breast cancer cohort via Cox proportional hazards testing. The top and bottom dendrograms show the results without and with adjusments of covariates for inflammation and proliferation. Blue and red are indicative of hazard ratio < 1 and hazard ratio > 1, respectively. All models included age and PAM50 subtype as covariates. B) Boxplots of gene set projection scores of selected REACTOME pathways, enriched in subgroups of immune cells. These pathways include: **PD-1 Signaling**, enriched in the *Trm All* subgroup, **Translation**, enriched in the *Translation*+ subgroup, **TNF Signaling**, enriched in the *CD4+ CCL5−* and *Treg TNFRSF4*+ subgroups, and **Cell Cycle**, enriched in the *CD8+ mit. Trm and Treg mit. Subgroups*. The center line, hinges, and whiskers indicate the median, interquartile range, and extreme values truncated at 1.5 * the interquartile range, respectively. C) Boxplots of markers constitutively regulated among selected *K2Taxonomer* subgroups. GZMB is upregulated in the *Trm All* subgroup. CCL5 and TNFRSF4 are up- and down-regulated, respectively, in the *CD4+ CCL5−* subgroup. TNFRSF4 is further up-regulated in the *Treg TNFRSF4+ subgroup,* while *RGS1*+ is up-regulated in the *Treg RGS1*+ subgroup. Finally, RPS27 is up-regulated in the *Translation*+ subgroup. The center line, hinges, and whiskers indicate the median, interquartile range, and extreme values truncated at 1.5 * the interquartile range, respectively. D) tSNE dimensionality reduction of the single-cell breast cancer immune cell data, indicating the cell subtype label assignment of every cell, as well as Z-scored expression of selected genes from C. E) 95% confidence intervals of hazard ratios from Cox proportional hazards testing of gene set projections of cellular subgroups on the *METABRIC* data set. *Covariates* shows the results of the survival model of sample-level inflammation and proliferation scores without a *K2Taxonomer* derived signature. Every other models shows the confidence interval of the subgroup-specific model without and with adjusting for inflammation and proliferation score, as well as the confidence intervals of inflammation and proliferation in the full model. All models included age and Pam50 breast cancer subtype as covariates. F) Scatterplot of the comparison of the expression of CCL5 and TNFRSF4 expression in the *METABRIC* dataset. G) 95% confidence intervals of hazard ratios from Cox proportional hazards testing of gene-level expression of CCL5 and TNFRSF4, modelled separately, *Sep.,* and combined in a single model, *Comb.*. These models also included age, Pam50 breast cancer subtype, as well as sample-level inflammation and proliferation score as covariates. H) Volcano plot of differential expression analysis of the *Translation*+ subgroup in scRNAseq data of individual genes in the REACTOME eukaryotic translation initiation gene set. An alternative coding of the *y-axis* indicating the absolute value of the test statistic is shown on the right side of the plot. The colors indicates the association of each gene with survival in the *METABRIC* data set. The names of genes, significantly associated with better survival (hazard ratios < 1, FDR < 0.1) are labelled. I) Comparison of the associations of expression of the REACTOME eukaryotic translation initiation gene set to survival in the *METABRIC* data set and the test statistics indicating up-regulation in the *Translation*+ subgroup. Genes that were included as top markers of the *Translation*+ subgroup are highlighted. The names of genes, significantly associated with better survival (hazard ratios < 1, FDR < 0.1) are labelled. The blue line indicates the linear fit of these two variables.

### Confounding effects of inflammation and proliferation on the association between tumor infiltrating cell activity and patient survival

To assess the clinical relevance of *K2Taxonomer* annotation of single-cell immune cell subgroups, we performed survival analysis, via Cox proportional hazards testing, modeling the relationship between *K2Taxonomer* subgroup gene signature scores and patient survival in the METABRIC breast cancer bulk gene expression data set.

For these models, we examined two possible sources of confounding factors. First, inflammation has a well-described paradoxical role in breast cancer progression^40^, such that the content of different subpopulations of lymphocytes has been associated with both better and worse prognosis^41^. Given the physiological similarities between different lymphocyte subtypes^42^, we hypothesized that expression patterns associated with tumor-promoting inflammation could mask those associated with tumor-suppressing TILs subsets. Second, we hypothesized that the signatures of the two mitotic T cell subgroups were similar enough to that of signatures of proliferative activity in non-immune tumor cells to result in a spurious association between T cell mitosis and worse prognosis. To assess and correct for these confounding effects, multivariate survival models were run without and with the inclusion of inflammation and proliferation scores as individual covariates. These patient-level scores were estimated by projecting published signatures of “bad” inflammation^43^ and proliferation^44^, each of which had been previously reported to be associated with poor prognosis in breast cancer.

The results of each of these analyses are summarized in Figure 5a. The full set of survival results for unadjusted and adjusted models, including the genes belonging to each subgroup signature are reported in Table S12. Controlling for inflammation and proliferation scores increased the overall significance of the association between subgroup-driven signatures of TILs and improved survival (hazard ratio < 1, FDR < 0.05). Furthermore, signatures of two cell subgroups, “CD8+ mit. Trm” and “Treg mit.”, characterized by increased cell cycle activity (Figure 5b), were associated with worse patient survival in models ignoring inflammation and proliferation scores, but were subsequently statistically insignificant in models including these covariates, likely reflecting the effect of confounding by proliferation activity (Figure 5a). This is further illustrated in Figure 5e, which shows the 95% confidence intervals of hazard ratios of “marginal” inflammation and proliferation models (left-most), as well as the confidence intervals of hazard ratios of select subgroups of cell subtypes, unadjusted and adjusted for inflammation and proliferation. Controlling for inflammation and proliferation allowed us to disentangle the contribution to survival of different components. For example, in the “CD8+ mit. Trm” subgroup, we observed that the “CD8+ mit. Trm” signature score was highly associated with worse patient survival in the unadjusted model, but the association became insignificant in the full model adjusted for proliferation and inflammation. On the other hand, there were instances where the hazard ratio achieved or improved significance (i.e., patient survival was significantly better) only after controlling for inflammation and proliferation in the full adjusted model, as observed in the “Trm All” subgroup and, to a lesser extent, in the “Translation+” subgroup (Figure 5e).

### High expression of TNFRSF4, a marker for Treg cell activity is associated with worse survival when adjusting for CCL5 expression

TNFRSF4 and CCL5 were found to be the top two markers constitutively up- and down-regulated, respectively, within Treg subgroups, with TNFRSF4 the top marker further discriminating between the two non-mitotic Treg subgroups, *Treg TNFRSF4*+ and *Treg RGS1*+ (Figure 5a, c, d). Furthermore, their expression was highly correlated in the METABRIC data set (rho = 0.66, p-value = 3.2e-245) (Figure 5f), supporting a pattern of co-expression within TIL microenvironments. To assess whether TNFRSF4 and CCL5 expression levels could serve as markers for immunosuppressive activity of Treg cells, we performed survival analysis of each gene modeled separately and in a combined model (Figure 5g). When modeled separately, the expression of TNFRSF4 is not associated with patient survival (P-Value = 0.32), while CCL5 is associated with better patient prognosis (P-Value = 1.94E-4). However, in the combined model both genes are associated with patient survival, with TNFRSF4 associated with worse patient survival (P-Value = 0.015).

Taken together these results indicate that, in bulk gene expression data, markers of Treg cell activity are highly correlated with markers of overall tumor immune infiltration, confounding associations between expression of these markers and patient survival.

### Up-regulation of specific translation genes characterizes a subgroup of TILs and is associated with better survival prognosis, independent of inflammation activity

The “Translation+” subgroup was a notable instance where the subgroup-specific signature projection was associated with better patient survival, regardless of adjustment for inflammation and proliferation (Figure 5a, e). To assess the extent to which up-regulation of translation-specific genes in this subgroup associated with better patient prognosis, we ran separate survival analysis for each of the 112 genes from the Reactome Eukaryotic Translation Initiation gene set, which were shared between the single-cell BRCA gene set and METABRIC data set. Of the 112 genes, 61 were up-regulated in the “Translation+” subgroup (FDR < 1E-5), including 26 genes within the top 50 marker “Translation+” subgroup signatures (Figure 5h, i). The full set of survival results for these 112 genes are shown in Table S13. The test statistics derived from single gene Cox proportional hazards models were negatively correlated with the corresponding genes’ test statistics of their up-regulation in the “Translation+” subgroup (rho = −0.23, p-value = 0.014) (Figure 5i). Furthermore, seven of the 112 genes were associated with better patient survival (FDR < 0.1). All of these genes were significantly up-regulated in the “Translation+” subgroup. Of these seven genes, RPL36A had the minimum “Translation+” subgroup associated test statistic (FDR = 1.34e-25) and two (RPS28 and RPS27) were members of the top 50 markers, comprising the “Translation+” subgroup signature. RPS27 was the top translation gene associated with the “Translation+” subgroup (FDR = 9.6e-246).

In summary, we employed *K2Taxonomer* to characterize the up-regulation of translational machinery as the dominant transcriptional program shared by a diverse subgroup of TILs. These findings informed additional analyses, which demonstrated that associations between expression of translational machinery genes and better patient survival were concordant with that of over-expression in TILs. Taken together these findings suggest that the up-regulation of specific translational machinery genes is widespread across TILs, serving as a predictor of the level of immune infiltration in breast cancer tissue from bulk gene expression data.

## Discussion

In this manuscript, we presented extensive assessment and practical applications of *K2Taxonomer*, a novel unsupervised recursive partitioning algorithm for taxonomy discovery in both bulk and single-cell high-throughput transcriptomic profiles. An important distinctive feature of the algorithm is that each partition is estimated based on a feature set selected to be most discriminatory within that partition, thus permitting the use of large sets of features to be used as input, without pre-filtering or dimensionality reduction approaches. Additionally, to minimize generalization error, each partition is based on an ensemble^45^ of partition estimates from repeated perturbations of the data. The adoption of an ensemble approach also makes it possible to compute a stability measure for each partition, which can be used to assess the robustness of each partition, as well as a stopping criterion for limiting the number of subgroup estimates. Finally, to facilitate comprehensive exploration of the results, as well as to share results for independent interrogation, the *K2Taxonomer* R package includes functionality to automatically generated interactive web-portals. One of these portals was utilized extensively to annotate subgroups as part of our analysis of breast cancer TILs in scRNAseq data, and is publicly available (https://montilab.bu.edu/k2BRCAtcell/).

As we have shown in its multiple applications, *K2Taxonomer* may be applied in a fully unsupervised mode to partition individual-level data, or it can take group-level labels as input to estimate inter-group relationships among the known groups. In the latter scenario, partition estimates are based on the constrained K-means algorithm^26^, which estimates clusters at the level of known group labels. This approach is perfectly suited to the downstream analysis of scRNAseq data, following the estimation of mutually exclusive cell types using scRNAseq clustering methods such as Seurat^14^ or Scran^15^. By preserving the single observation information within each group, and by thus being able to tailor the feature set to each of the groups, we expect our approach to outperform methods in which group-level information is summarized into single statistical measures. This conclusion is supported by our simulation analysis, where *K2Taxonomer* was shown to significantly outperform Ward’s agglomerative method based on group-level test statistics. Even when adopted for observation-level analysis, where inference was performed on the full set of individual observations, *K2Taxonomer* was still shown to significantly outperform standard agglomerative methods, on both simulated and real data, although not to as large an extent.

In our analysis of healthy airway cell types’ annotation^28^, we employed *K2Taxonomer* to (re)discover subgroups of cell types characterized by shared lineage. Remarkably, our analysis accurately recapitulated the known taxonomic structure relating the different cell types to an extent not matched by the other methods evaluated. This example illustrates a prototypical use of the tool: in those cases where a data set and its associated cell type estimations are publicly available, *K2Taxonomer* facilitates their immediate repurposing for additional insight and discovery.

It is important to emphasize that in many data sets continuous lineage trajectories are non-existent or obscured by phenotype-driven inter-group transcriptional relationships. While *K2Taxonomer* cannot identify precursor relationships between cell types, the strategy of pairing recursive partitioning with local feature selection allows the discernment of relative relationships between groups rather than “all-or-nothing” connections as in graph-based trajectory models. The advantages of this strategy are exemplified by the analysis of the healthy airway, in which the data included samples of multiple individuals and airway locations. *PAGA* analysis of these data produced highly-connected graphs with no decipherable trajectories or subgroups beyond that of disconnected subgraphs. On the other hand, *K2Taxonomer* recapitulated clear lineage-driven hierarchies of airway cell subgroups. While the subgrouping of endothelial cells as a subset of immune cells does not reflect the expected hierarchy, *K2Taxonomer* generated this model without parameterization. The “best” *PAGA* model had the unfair advantage of having been chosen from a set of distinct models generated from combinations of feature selection and dimensionality reduction parameters. In practice, the a priori choice of the optimal values of these parameters is challenging.

Our extensive analysis of single-cell data from breast cancer TILs showcased the incorporation of *K2Taxonomer* in an advanced in-silico study that yielded significant novel insights. In contrast to bulk gene expression, which captures average expression across all cells, identifying dominant transcriptional programs driving phenotypic similarities between subgroups of cell populations offers additional insights to deconvolute the cellular microenvironment of these samples beyond their individual transcriptional signatures. Molecular convergence of cells of disparate lineages is exemplified by subpopulations of CD8+ and CD4+ T cells, each of which exists in various functional states as naïve, effector, and memory subpopulations^46^. Importantly, our K2T-based analysis showed that concordant subpopulations of CD8+ and CD4+ T cells share transcriptional signatures that may outweigh those arising from their shared lineage. For example, both CD8+ Trm and CD4+ Trm cells have been reported to express surface molecules, CD69 and CD11a^47^. Concordantly, CD8+ Trm and CD4+ Trm cells were segregated into a common subgroup by K2T, demonstrating the relative dominance of their shared transcriptional activity. Projection of the expression signature of Trm cell subgroups was associated with better survival in the METABRIC data set. Past studies focusing on CD8+ Trm cell markers have reported similar findings^29,48^.

Unlike Trm cells, the presence of immune-suppressing Treg cells in the micro-environment has been associated with poor prognosis in breast cancer^49–51^. After identifying TNFRSF4 as heterogeneously expressed across the Treg cell subgroup, we showed that TNFRSF4 expression was associated with worse patient survival in the METABRIC data set when adjusted for CCL5 expression, which was down-regulated among all Treg cells. This supports previous findings that TNFRSF4, also known as OX40, is a marker of high Treg cell immunosuppressive activity^52,53^. The high level of co-expression of TNFRSF4 and CCL5 in the METABRIC data set suggests that either gene is associated with immune infiltration in breast cancer tumors. Additionally, this provides a resolution as to why projections of the signature of the Treg cell subgroup were associated with better patient survival, while the signature of the Treg cell subset, characterized by high TNFRSF4 expression, was not. These results are consistent with previous studies establishing the ratio between Treg and CD8+ T cell abundance as a prognostic marker of breast cancer that reflects immune inhibitory function of Treg cells^54^. Moreover, these results strongly suggest a prognostic value for markers that capture the degree of immune inhibitory activity of Treg cell populations.

Finally, *K2Taxonomer* identified a diverse subgroup of breast cancer TILs characterized by consistent up-regulation of translational genes. Increased ribosomal biogenesis has been previously implicated in increased tumorigenesis^55–58^, but has only recently been implicated in T cell activation^30^ and expansion^31^. Unlike the majority of other subgroups, the signature of the T cell subgroup overexpressing translational machinery genes was associated with better patient survival in METABRIC patients regardless of adjustments for inflammation^43^ and proliferation^44^ signatures. Furthermore, the association of the expression of specific translational genes with better patient survival was significantly correlated to their overexpression in this T cell subgroup. These results suggest that overexpression of these T cell-specific translational genes is not masked by tumor-specific gene expression and is therefore indicative of CD4+ and CD8+ T cell tumor infiltration.

In summary, *K2Taxonomer* demonstrated a remarkable ability to discover biologically relevant taxonomies when applied to the analysis of both bulk gene expression and scRNAseq data and to outperform standard agglomerative methods. In multiple practical applications, we showcased the versatility of *K2Taxonomer* to analyze scRNAseq data toward the characterization of genes and pathways distinguishing specific subgroups, thereby generating hypotheses that were then in-silico validated in independent bulk gene expression data. As noted, while we here focused on the analysis of transcriptomics data, the proposed approach is equally applicable to other bulk and single-cell ‘omics’ data, such as those generated by high-throughput proteomics and metabolomics assays.

## Methods

### K2Taxonomer Algorithm Overview

*K2Taxonomer* implements a recursive partitioning algorithm that takes as input either a set of individual observations or a set of sample groups and returns a top-down hierarchical taxonomy of those samples or groups (Figure 1). To achieve robust model estimation, each partition is defined based on the aggregation of repeated partition estimations from distinct perturbations of the original set. Each of these partition estimates is created in three steps. First, a perturbation-specific data set is generated by bootstrapping features, i.e., sampling features from the original data set with replacement. Next, this perturbation-specific data set undergoes variability-based feature selection filtering. Finally, a K=2 clustering algorithm is run, producing a perturbation-specific partition estimate. These three steps are repeated, generating a set of perturbation-specific partition estimates, which are aggregated into a cosine similarity matrix (see Supplementary Methods). The aggregate partition is then estimated based on a K=2 tree cut following hierarchical clustering of the transformed cosine similarity matrix into a distance matrix, calculated as 1 - cosine similarity, with a user-specified agglomeration method.

The current implementation of *K2Taxonomer* includes many options for parametrizing steps in this procedure. By default, *K2Taxonomer* performs these perturbation-specific partition estimates via agglomerative clustering of the Euclidean distance matrix of the bootstrapped data set. Additionally, the *K2Taxonomer* package includes functionalities for performing group-level recursive partitions, i.e., partitioning data sets where observations have a priori-assigned group labels, whereby the objective of the *K2Taxonomer* procedure is to identify intermediate relationships between these groups. This functionality was specifically incorporated to enable partitioning and annotations of cell types estimated by scRNAseq clustering algorithms, but it is applicable to any data set with group-level labels. Finally, to allow for further customization of analyses, the *K2Taxonomer* R package permits the use of user-specified functions for performing perturbation-specific partition estimates.

### K2Taxonomer R Package Functionalities

In addition to running the recursive partitioning algorithm, the *K2Taxonomer* R package provides functionalities for comprehensive annotation of the estimated subgroups, via subgroup-level statistical analyses, including: differential analysis, gene set enrichment analysis, and phenotypic variable testing. Differential analysis of gene expression is carried out using the *limma* R package, which is well-suited to the analysis of normally distributed data such as microarray gene expression, as well as log-transformed and normalized RNAseq data^59^. Gene set enrichment analysis is carried out on a set of user-provided gene sets and implemented in two ways: over-representation analysis based on a hypergeometric test, and differential analysis of single-sample gene set projections scores based on the *GSVA* R package^60^. Finally, phenotypic variable testing is carried out on user-provided variables labeling individual observations or groups, supporting both continuous and categorical variables. Testing of association between continuous variable and taxonomy subgroups can be performed based on the parametric Student’s t-test or the nonparametric Wilcoxon rank-sum test, while categorical testing is carried out using Fisher’s exact test. All subgroup-level statistical analyses are corrected for multiple hypothesis testing based on the FDR procedure^61^. The full set of results are compiled into an interactive-web portal for exploration and visualization. Differential analysis comparisons are carried out at the partition-level, i.e., comparing only the two subgroups at a particular node. However, the web portal includes functionality for performing post-hoc differential analysis of any combination of user-selected subgroups.

### Statistical analysis

The implementation of *K2Taxonomer* for this manuscript was run with *R (v3.6.0)*, *limma (v3.42.2)*, and *GSVA (v1.34.0)*. All p-values reported in this manuscript are two-sided.

### Simulation-based performance assessment

The performance of *K2Taxonomer* was assessed in comparison to Ward’s agglomerative method for recapitulating induced hierarchical structure in simulated data. See supplementary methods for a comprehensive description of the strategy implemented for data generation and performance assessment for observation- and group-level analyses.

### METABRIC breast cancer primary tumor bulk gene expression processing

METABRIC breast cancer primary tumor Illumina HT-12 v3 microarray bulk gene expression data was obtained from the *CBioPortal*, https://www.cbioportal.org/datasets^62,63^. The data set includes normalized expression values for 24,360 genes and Pam50 cancer subtype estimations for individual primary breast cancer tumor samples across 1,974 female patients. Additional clinical variables considered for this analysis, included: patient age at diagnosis, survival status, ER-status, PR-status, and HER2-status. The distribution of these variables across patients is summarized in Table S4.

### TCGA breast cancer primary tumor bulk gene expression processing

The *cancer genome atlas (TCGA)* breast cancer (BRCA) primary tumor bulk RNAseq data was obtained from *Genomic Data Commons (GDC)*, https://gdc.cancer.gov/access-data/^64^. The data set includes raw gene expression counts for 36,812 genes and Pam50 cancer subtype estimations for individual primary breast cancer tumor samples across 973 female patients. Additional utilized clinical variables, included: patient age at diagnosis, ER-status, PR-status, and HER2-status. The distribution of these variables is summarized in Table S5.

Raw counts were normalized by the trimmed mean of M-values (TMM) method and log-normalized using *edgeR (v3.28.1)* R package^65^ and genes with fewer than 2 reads in more than 90% of samples were removed, resulting in 25,729 genes in the processed data set.

### Performance assessment using breast cancer primary tumor bulk gene expression data

*K2Taxonomer* was evaluated for its ability to recover the Pam50 subtypes [REF], as well ER-, PR-, and HER2-status, and the aggregate three-gene genotype of ER-, PR-, and HER2-status, in the *TCGA* and *METABRIC* breast cancer datasets, independently. *K2Taxonomer* was also compared to two agglomerative clustering algorithms, Ward’s and average. These specific methods were chosen because they have been previously shown to outperform other common agglomerative methods^22,66^. Given the sensitivity of hierarchical clustering to the level of feature filtering, analyses included individual runs on four filtered data subsets of the total number of features: 100%, 25%, 10%, and 5%, while *K2Taxonomer* was only run on the full set of 100% of the total number of features. This should be kept in mind when comparing performances, since the best-performing pre-filtering level is not known a priori, and it is in general dataset dependent. For every pre-filtering level, the median absolute deviation (MAD) score was used for feature selection, and Euclidean distance was used to estimate observation-level distance. Performance was assessed as the entropy of each of the phenotypes (e.g., PAM50 labels), induced by the inferred sample sub-grouping, with lower entropy indicating “purer” subgroups, hence better performance^67^. The different methods were evaluated and compared by the relative decrease in entropy as the number of mutually exclusive clusters, K, increased from 2 to 8 based on tree cuts of the dendrograms produced by each model.

### Healthy airway tissue scRNAseq gene expression analysis

Publicly available scRNAseq data of normalized, batch corrected, and log-transformed gene expression estimates from airway tissue of healthy subjects were obtained from the *UCSC Cell Browser* portal, published as a supplement to the original manuscript for which these data were used, https://www.genomique.eu/cellbrowser/HCA/?ds=HCA_airway_epithelium^28^. This data set includes expression estimates for 18,417 genes and 77,969 individual cells from 35 samples across 10 subjects. Multiple samples taken from individual subjects were collected from distinct locations of the human airway including: nasal biopsies, nasal brushings, tracheal biopsies, intermediate bronchial biopsies, and distal brushings. Also, this data set included cell type estimations for each of the 77,969 cells and comprised 28 estimated cell types in total. The methods of data processing, as well as distributions of subject-level sample identities and estimated cell types, can be found in the original publication^28^.

*K2Taxonomer* partitioning of these 28 estimated cell types was evaluated against the known relationships among the included cell types and was compared to the partitioning obtained by two agglomerative methods, Ward’s and average, as well as PAGA, a graph-estimation and trajectory inference aglorithm^20^. For agglomerative methods, cell type-level data processing and feature selection were performed consistent with the results of group-level analysis of simulated data (See supplementary methods). Analyses with *PAGA* were carried out using the *scanpy* (v1.6.0) python package. As with agglomerative methods, PAGA was run on F-statistic-based pre-filtered feature sets at different percentages of the full 18,417 genes: 100%, 25%, 10%, and 5%. Following component-based dimensionality reduction of these feature sets, the neighborhood graphs were estimated using “pp.neighbors()” for three numbers of principal components: 10, 15, and 20, such that a total of 12 unique *PAGA* models were estimated. Finally, the *PAGA* algorithm was run, setting the “group” argument of “tl.paga()” as the cell type labels.

### Breast cancer immune cell scRNAseq gene expression analysis

Publicly available scRNAseq gene expression of raw counts from immunocytes of two TNBC patients was obtained from *GEO*, accession number GSE110938^29^. The data was processed in accordance with the original manuscript^29^, recapitulating the reported 5,759 individual cells, 4,844 and 915 from either sample, with 15,623 genes passing QC criteria, selection of 1,675 highly variable genes, and 10 latent variables estimated by *ZINB-WaVE* (v1.8.0)^68^. To enable exploration of the data at finer resolution, clustering of the latent variables with Seurat (v1.3.4) was modified by setting the “resolution” argument of “FindClusters()” to 1.1, rather than the default, 0.8^69^. This resulted in 13 estimated cell clusters. Of the 10 cell clusters reported in the original manuscript, two cell clusters, “CD4+ FOXP3+” and “CD4+ IL7R+”, were further split into three and two individual clusters, respectively (Table S8).

*K2Taxonomer* partitioning of these 13 estimated cell subtypes was performed on the normalized count matrix estimated by *ZINB-WaVE*. According to the developers, *ZINB-WaVE*, normalized count estimates are not recommended for differential analysis^68^, hence differential analysis was performed based on drop-out imputed and batch-corrected normalized counts estimated using the *bayNorm (v1.4.14)* R package^70^. Pathway-level analysis was carried out using Reactome gene sets downloaded from *mSigDB (v7.0)*^71^. Signatures of up-regulated genes were derived from each subgroup based on their FDR corrected p-value (FDR < 1e-10) and minimum subgroup-specific expression, (mean[log2 counts] > 0.5), then restricted to a maximum of 50 genes.

To validate the clinical relevance of signatures of tumor infiltrating lymphocytes (TILs) derived by *K2Taxonomer* we performed survival analysis based on gene signature projection scores, as well as on selected genes in the *METABRIC* breast cancer primary tumor gene expression data set. Gene set projection was carried out using *GSVA*^60^. Multivariate Survival analysis was performed using Cox proportional hazards tests. All models included age and Pam50 subtype as covariates. To account for possible confounding effects of inflammation and proliferation, we generated separate patient-level activity scores for each, using a gene set projections of published signatures of deleterious breast cancer inflammation markers^43^ and breast cancer proliferation^44^ (Table S11).

## Acknowledgments

The results shown here are in part based upon data generated by the TCGA Research Network: https://www.cancer.gov/tcga. We would like to thank Bob Varelas and Gerald Denis for useful feedback on the Breast Cancer TIL case study.

## Data Availability

The data that support these findings are publicly available and were accessed from several repositories. The bulk microarray gene expression and subject data from the METABRIC breast cancer study is available through the *cBioPortal*, https://www.cbioportal.org/study/summary?id=brca_metabric. The bulk RNAseq gene expression and subject data from the TCGA BRCA project is available through the *Genomic Data Commons*, https://gdc.cancer.gov/access-data/. The single-cell RNAseq gene expression and subject data of healthy airway tissue is available through the *UCSC Cell Browser page*, https://www.genomique.eu/cellbrowser/HCA/?ds=HCA_airway_epithelium. Finally, the single-cell RNAseq gene expression and subject data of breast cancer immune cells is available in the *Gene Expression Omnibus*, <u>GSE110938</u>.

## Supplementary Figures

**Figure S1:**
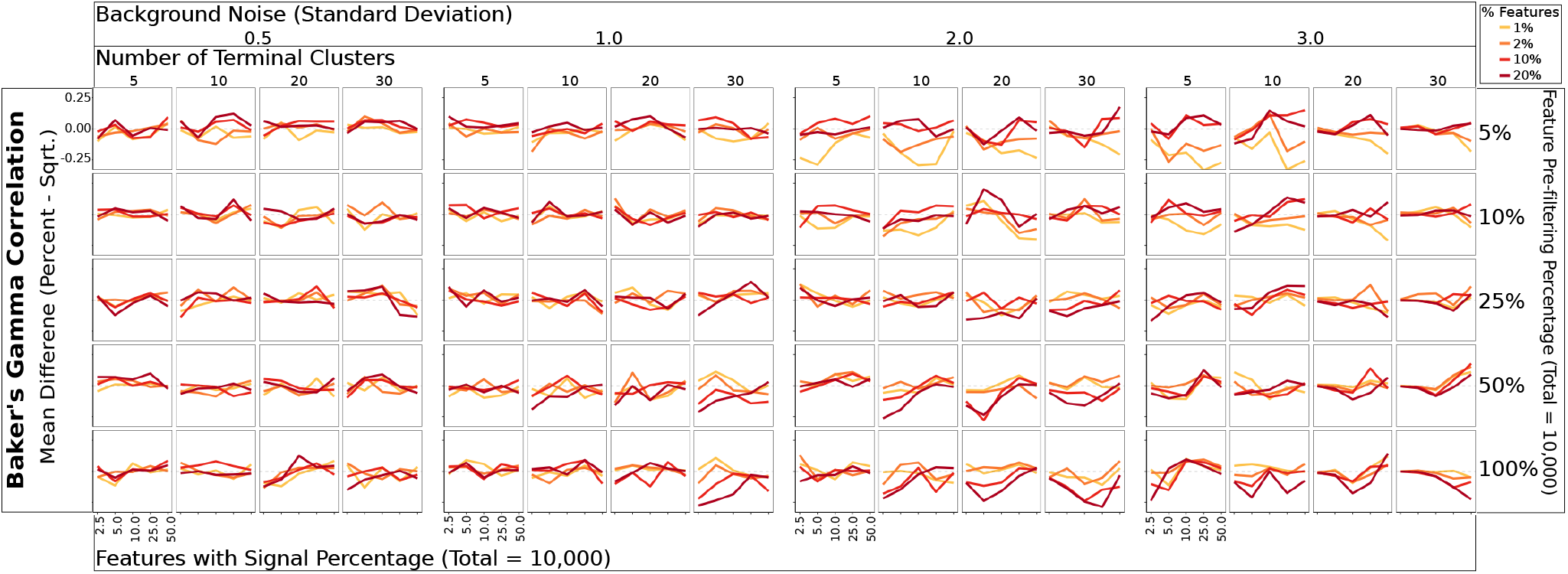
Simulation-based performance assessment of running *K2 Taxonomer* using different partition-specific feature subset sizes of the full data set. Difference of mean Baker’s gamma correlation estimates measuring the similarity of *K2 Taxonomer* estimates to the true hierarchy from which the simulated data was generated between. Each line shows the difference between a set percentage of the total number of features and the square root of the total number of features. Each combination of parameters was simulated 25 times.

**Figure S2:**
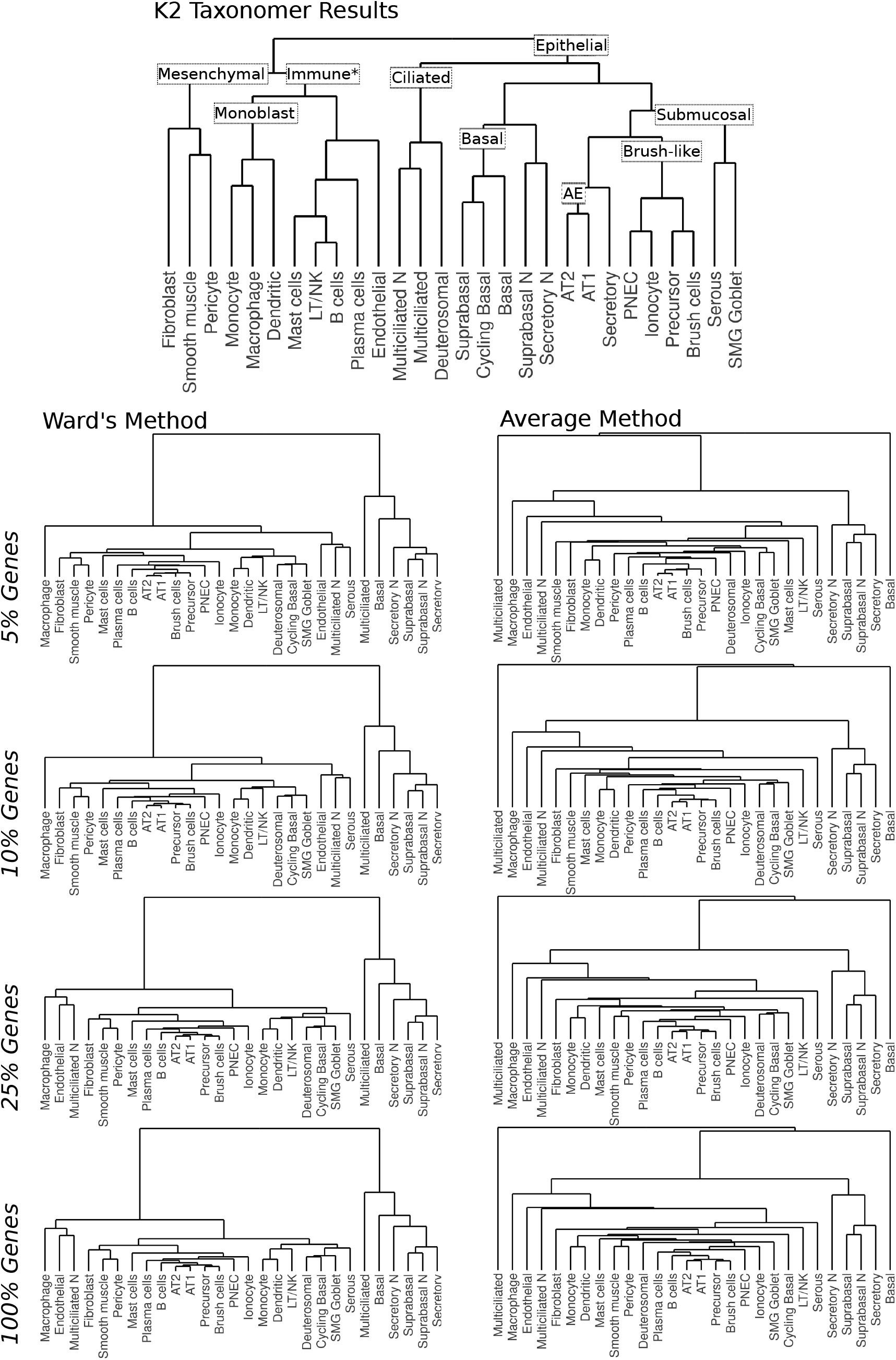
Additional subgrouping results of healthy airway cell types from scRNAseq data. Ward’s (left) and average (right) agglomerative clustering results for analyses performed on different subsets of the total number of genes.

**Figure S3:**
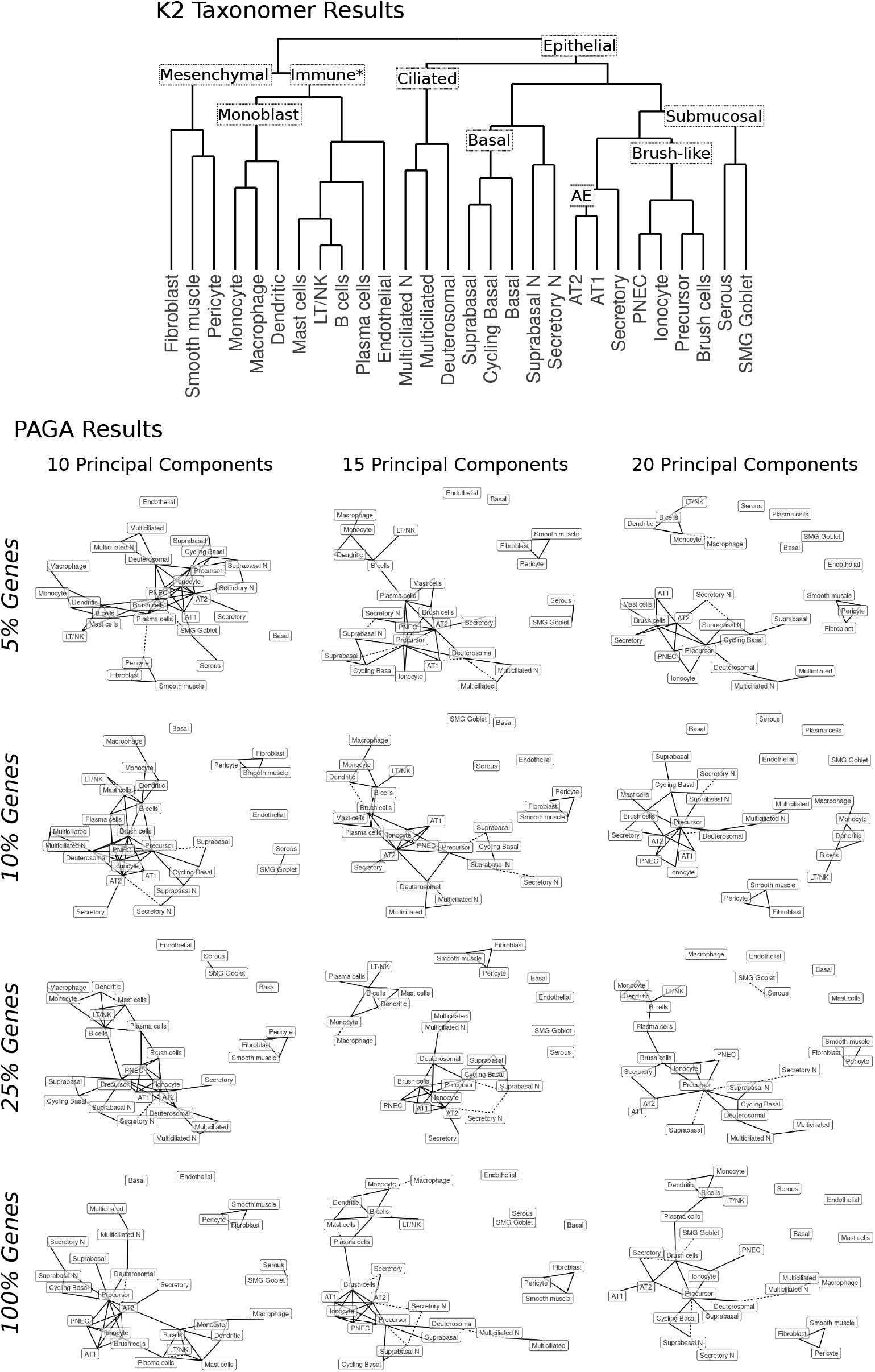
Additional PAGA graph-based trajectory results of healthy airway cell types from scRNAseq data. PAGA graph-based trajectory results for analyses performed on different combinations of numbers of principal components and subsets of the total number of genes.

## Supplementary Methods

### K2Taxonomer feature filtering

A distinguishing property of *K2Taxonomer* when compared to other methods, such as traditional agglomerative hierarchical cluster or trajectory inference^1^is the manner in which feature selection is implemented. Even in large studies of high-throughput data sets, the number of features is typically much larger than the number of observations. This generally requires filtering the data set prior to modeling in order to reduce variance and computational expense of model fit. One way to do this is through feature selection, in which features suspected to contain more information about the relationship between observations are chosen for down-stream analysis. For unsupervised learning, relative information estimation is commonly calculated via variability-based metrics. Assuming the amount of noise is consistent across features, these metrics will capture the relative magnitude of the signal of individual features. Two common choices are standard deviation (SD) and median absolute deviation (MAD), of which the former is more statistically efficient with a small sample size and the latter is more robust to outliers^2^. Implementation of these feature selection techniques prior to modeling may be problematic when learning hierarchical models. The magnitude of variability-based metrics is influenced by the frequency of observations for which the signal-to-noise ratio is higher, such that the subset of features is more likely to capture broader relationships between larger sets of observations and less likely to capture relationships between smaller sets of observations. This can obscure important relationships within smaller sets of observations, as when evaluating a sub-group of samples in a hierarchical procedure. In addition, an appropriate choice of the number features to use for modeling is difficult to determine a priori and may be obscured by many factors, including: the number of subgroups, number of observations belonging to each subgroup, and the number of features distinguishing individual subgroups.

To overcome these challenges, *K2Taxonomer* produces a model fit for each partition independently, such that feature selection is only performed within the subgroup of observations being evaluated at a given step. In particular, at each recursive step, the objective of partition estimation is to split the data based only on the dominant relationship between two subgroups. Since the selected features need only capture one relationship, a much smaller subset of features will be sufficient to discover this partition. By default, *K2Taxonomer* uses the square root of the total number of features, which is used in a related albeit supervised learning method, random forests^3^. In doing so, the percentage of filtered features is dependent on the total number of features. For example, if the data set consists of 1,000 or 10,000 features, *K2Taxonomer* will estimate partitions using 3.2% or 1.0% of the total number of features, respectively. The appropriateness of using the square root of the total number of features against fixed percentages is later assessed with simulation-based testing.

The *K2Taxonomer* package includes options to perform both SD and MAD based feature selection. In the case of group-level analysis, *K2Taxonomer* can perform F-statistic based feature selection based on the proportion of between-group variability and within-group variability implemented by the *limma* R package^4^.

### K2Taxonomer data partitioning

To estimate each partition, *K2Taxonomer*, performs feature-level bootstrap aggregation, similar to that of consensus clustering^5^. More specifically, each data partition represents the aggregation of a set of partitions estimated from perturbations of the original data set in which features have been sampled with replacement. Feature selection and K=2 clustering are independently performed within each perturbation-specific data set. The final partition estimate is calculated by aggregating the set of perturbation-specific partitions into a *cosine similarity* matrix (defined below), which further undergoes hierarchical clustering, followed by a K=2 tree cut.

*K2Taxonomer* package implements separate clustering methods tailored to analysis of either observation-level and group-level data input. For observation-level data, the perturbation-specific partitions are estimated via hierarchical clustering of the Euclidean distance matrix, followed by a K=2 tree cut. By default, Ward’s agglomerative method is performed at this step because it has been shown to generally perform well compared to other hierarchical methods^6^. For group-level data, perturbation-specific partitions are estimated via constrained K-means clustering^7^. This algorithm performs semi-supervised clustering, in which group-level information is included as a pairwise “must-link” constraint, preserving relationships between observations from the same group.

To assess the robustness of the partitioning of the aggregated results, hereby referred to as partition stability, as well as to facilitate interpretability, a cosine similarity matrix is computed, with each pairwise cosine similarity measurement functionally equivalent to the Pearson correlation of standardized variables.

Let an “item” denote a single observation or group, depending on whether observation- or group-level analysis is being performed, respectively. The cosine similarity of two items is a measure proportional to the number of times across perturbation iterations that the two items are assigned to the same group in the perturbation-specific dichotomous partitions. It takes its maximum/minimum value when the two items are always/never assigned to the same group.

If we represent with “−1” and “1” the assignments of an item to one or the other group in a dichotomous partition, we can then represent, and compare, the complete set of assignments of any two items across *p* perturbation-specific partitions as the vectors

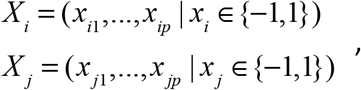

where *X*_*i*_ and *X*_*j*_ represent the *i*^*th*^ and *j*^*th*^ item, respectively.

We can then define the cosine similarity of *X*_*i*_ and *X*_*j*_ as

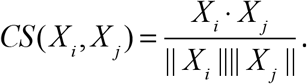

where *X*_*i*_ · *X*_*j*_ represents the dot product and || *X*_*i*_||||*X*_*j*_|| represents the product of the Euclidean norms of *X*_*i*_ and *X*_*j*_. Next, we prove the equivalence of the cosine similarity and Pearson correlation of two assignment vectors. The cosine similarity can be rewritten as follows:

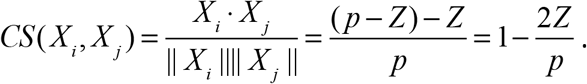

In the above derivation, since *X*_*i*_ and *X*_*j*_ only take values in {−1,1}, their dot product, *X* ·*Y*, is equal to the difference between the number of iterations, *Z*, the two items are assigned to the same group, and the number of iterations, *p-Z*, the two items are assigned to different groups. Furthermore, the product of the Euclidean norms of *X*_*i*_ and *X*_*j*_, || *X*_*i*_ |||| *X*_*j*_ ||, is equal to *p*.

Similarly, taking advantage of the relationship between Pearson correlation of standardized variables, *r*(), and Euclidean distance, *d*(), we have

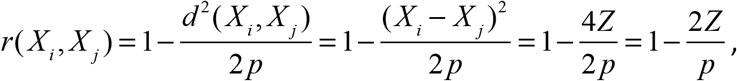

where we used the fact that the squared Euclidean distance of *X*_*i*_ and *X*_*j*_, *d*^2^ (*X*_*i*_, *X*_*j*_), is equal to (*X*_*i*_ − *X*_*j*_)^2^. Furthermore, for *X*_*i*_ − *X*_*j*_, the difference between mismatched adjacent elements is 2 and the difference between matched adjacent elements is 0. Therefore, (*X*_*i*_ − *X*_*j*_)^2^ = 4*Z*.

The function, 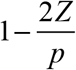, is the Hamann similarity index^8^. Consistent with Pearson correlation, the range of possible values for the Hamann similarity is between 1 and −1; these extremes occur if *Z* is equal to *p* and 0, respectively, indicating that *X*_*i*_ and *X*_*j*_ are either identical or fully dissimilar. Furthermore, if elements of *X*_*i*_ and *X*_*j*_ share 50% of their matching assignments, then 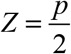 and the Hamann similarity is equal to 0, indicating a lack of a relationship between their perturbation-specific partition estimates. This is not true for the related phi similarity^9^, another correlation metric for dichotomous variables, the calculation of which includes adjustment for the marginal distribution of *X*_*i*_ and *X*_*j*_. In this case, the marginal distribution of *X*_*i*_ and *X*_*j*_ is irrelevant because the elements of *X*_*i*_ and *X*_*j*_ are only meaningful in relation to their matching assignments.

### K2Taxonomer Partition Stability

To assess the robustness of partition estimates, indicating the consistency of the perturbation-specific partition results, we developed a partition stability metric, which is calculated using the eigen-decomposition of the matrix of pairwise cosine similarities, *Q*, of dimension, *N*, the number of items. The eigen-decomposition of *Q* satisfies

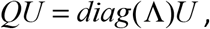

where *U* is the matrix of eigenvectors corresponding with Λ, the vector of rank-ordered eigenvalues

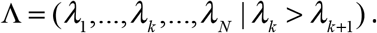

Each eigenvalue is proportional to the “variance explained” by each eigenvector, such that the cumulative sum of variance explained by the first *k* eigenvectors, *v*_*k*_, is given by

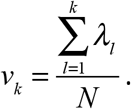

In this context, the variance explained by eigenvectors captures the consistency with which pairs of items received the same or different assignments across perturbation-specific partitions. Therefore, we can summarize this consistency by evaluating the difference between the variance explained by the eigenvectors of the estimated cosine matrix and the variance explained by these eigenvectors *if* there was no consistency across perturbation-specific partition assignments. We denote this deviation as the partition stability, *PS*, calculated as the maximum difference between *v*_*k*_ and 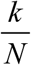, the null value corresponding to all items being linearly independent,

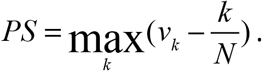

The possible values for the partition stability range from 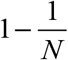 and 0, with the former representing the case in which *λ*_1_ = *N*, where every perturbation-specific partition is identical, i.e., all values of the cosine matrix are either −1 or 1. Conversely, a partition stability of 0 represents the case when the perturbation-specific partition assignments are random, i.e., all values of the cosine matrix are close to 0. The maximum value for a given partition is dependent on the number of items in the partition, approaching 1 when *N* is large, and equal to 0.5 when *N* = 2. Using the *K2Taxonomer* package, partition stability can be used to set stopping criteria for creating new partitions, thereby serving as a way to control the number of terminal subgroups without prior knowledge.

Finally, partition stability is used as a heuristic for calculating branch heights in dendrogram creation of the *K2Taxonomer* output. For a series of *m* partitions resulting in a given partition, *z*_*m*_, the branch height, *h*_*m*_, is calculated as

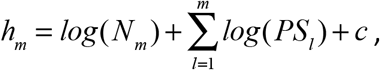

where *c* is a constant added to ensure that the minimum height for a node is equal to 1.

### Simulated data generation

*K2Taxonomer*’s performance was assessed in terms of its capability to recapitulate induced hierarchical structure in simulated data, and it was further compared to Ward’s agglomerative method as a term of reference. Hierarchically structured data was generated by assigning a mutually exclusive set of *C* labels to *N* observations. To define hierarchical relationships between labels, the set of *C* labels was recursively subdivided into intermediate subgroups until the final subgroups contained only a single label. To represent expected hierarchical structures in real data, we allowed for more than two subgroups to be created at each subdivision. As such, neither *K2Taxonomer* nor agglomerative methods are expected to recapitulate this structure exactly.

After simulating the hierarchical relationships between labels, data was generated as follows. First, a set of 10,000 features was generated, each from a normal distribution, with mean, 0, and standard deviation, BN, denoting the background noise in the data. Second, for each subgroup of labels, we assigned a random set of features for which to add a signal. For each feature assigned to a specific subgroup of labels, the value of the added signal was sampled from a normal distribution with mean, 0, and standard deviation, 2. To ensure that the value of the added signal was the same across all subgroup-specific observations, a feature was only allowed to be assigned to an individual label once. Considering that in real data we expect only a subset of features in a data set contains information of its subgroup structure, prior to assigning random sets of features to each subgroup, the full set of features was subsetted by a given percentage, DS, of the total.

### Simulated data performance assessment

Following the generation of each simulated data set, *K2Taxonomer* and Ward’s method were run. The performance of each method was assessed by the relative similarity of the learned structure to the known hierarchy from which the data was generated using Baker’s gamma correlation^10^. Baker’s gamma correlation is a measure of Spearman correlation between two similarity matrices, where each similarity matrix is calculated from the number of shared partitions between each pair observations in a given hierarchy. Baker’s gamma correlation ranges between −1 and 1, with 1 representing the case when two dendrograms induce identical hierarchies. This metric was chosen over *cophenetic distance* because *cophenetic* pairwise similarities are calculated using branch heights and branch height is not applicable to the known hierarchy.

### Observation-level analysis of simulated data

For observation-level analysis, each simulated data set included N=300 observations. We evaluated the following data generation parameters

- Number of labels (C): 5, 10, 20, 30
- Background Noise (BN): 0.5, 1, 2, 3
- Percent features with signal (DS): 2.5, 5, 10, 25, 50.

Especially when the value of DS is small, Ward’s method is not expected to perform well when run on the full set of data. Therefore, Ward’s method was evaluated on simulated data with the following variability-based prefiltering percentage levels, PF, of the data:

- Pre-filtering percentage level (PF): 5, 10, 25, 5, 1.

On the other hand, *K2Taxonomer* was always run on the square root of the total number of features. It should be emphasized that this design does not allow for an unbiased comparison of K2T and Ward, and it favors the latter since in real settings we would not know the optimal PF to be used.

Standard deviation was used for variability-based feature selection, including partition-specific feature selection by *K2Taxonomer*. For each combination of these parameters, we generated 25 simulated data sets and ran *K2Taxonomer* and Ward’s method. For each combination of parameters, the statistical significance of the difference between the two methods’ distributions of Baker’s gamma correlation estimates was tested using Wilcox’s Rank Sum tests. The resulting p-values from 400 combinations of parameters were corrected for multiple hypotheses testing using the FDR procedure^11^.

As noted, for each of these comparisons, we ran *K2Taxonomer* using the square root of the total number of features as the parameter for partition-specific feature filtering. To assess the validity of using this as the default, we performed additional simulations and compared *K2Taxonomer* performance at different partition-specific feature filtering levels, based on the set of percentages of the total number of features: 1%, 2%, 10%, and 20%. These were performed using the same combinations of parameters as the previous analysis and included 25 repetitions.

### Group-level analysis of simulated data

For group-level analysis, each simulated data set included N=1000 observations. For data generation, we tested the following sets of data generation parameters

- Number of labels (C): 10, 25, 20, 30
- Background Noise (BN): 0.5, 1, 2, 3
- Percent features with signal (DS): 2.5, 5, 10, 25, 50.

As with observation-level analysis, we performed analyses on simulated data with the following variability-based prefiltering percentage levels, PF, of the data:

- Pre-filtering percentage level (PF): 5, 10, 25, 5, 1.

Linear model-based F-statistics were used for all variability-based feature selection scoring, including partition-specific feature selection by *K2Taxonomer*.

For running each method, the groups were defined by the labels to which each of the observations were assigned. Unlike *K2Taxonomer*, agglomerative methods are not devised to utilize group-level labels for unsupervised learning tasks. Therefore, Ward’s method was applied using the Z-score mean value of each group, generated by the same linear model used for feature selection. Comparisons between these methods were performed in the same manner as observation-level analysis.

